# Scalable hierarchical clustering by composition rank vector encoding and tree structure

**DOI:** 10.1101/2020.04.12.038026

**Authors:** Xiao Lai, Pu Tian

## Abstract

Supervised machine learning, especially deep learning based on a wide variety of neural network architectures, have contributed tremendously to fields such as marketing, computer vision and natural language processing. However, development of unsupervised machine learning algorithms has been a bottleneck of artificial intelligence. Clustering is a fundamental unsupervised task in many different subjects. Unfortunately, no present algorithm is satisfactory for clustering of high dimensional data with strong nonlinear correlations. In this work, we propose a simple and highly efficient hierarchical clustering algorithm based on encoding by composition rank vectors and tree structure, and demonstrate its utility with clustering of protein structural domains. No record comparison, which is an expensive and essential common step to all present clustering algorithms, is involved. Consequently, it achieves linear time and space computational complexity hierarchical clustering, thus applicable to arbitrarily large datasets. The key factor in this algorithm is definition of composition, which is dependent upon physical nature of target data and therefore need to be constructed case by case. Nonetheless, the algorithm is general and applicable to any high dimensional data with strong nonlinear correlations. We hope this algorithm to inspire a rich research field of encoding based clustering well beyond composition rank vector trees.

## Introduction

### Present clustering algorithms and challenges

Clustering is a ubiquitous task in many subjects such as chemistry, physics, biology and machine learning, and existing algorithms are as expectedly quite diverse. These algorithms may be classified as partition, hierarchical algorithms and various forms of their hybrids. K-means^1^ is arguably the simplest and most widely utilized partition clustering algorithm. Many other clustering algorithms (e.g. meanshift,^2^ dbscan,^3^ cure,^4^ affinity propagation,^5^ spectral clustering,^6,7^ …), while have various assumptions/complexities/advantages, share the distance/similarity metric calculation for pairwise comparison between records or their clusters, or that between records and specified points, which engenders significant computational cost regardless of specifics. General distance metrics (e.g. Euclidean, Manhattan distances) provide weak differentiating power in high dimensional space. To tackle this issue, various dimensionality reduction methods are utilized with principal component analysis (PCA)^8^ being the most populous. However, PCA requires expensive matrix diagonalization and is unable to deal with nonlinear correlations. More complexities (e.g. kernels) introduced for treatment of nonlinearity bring in further uncertainties and difficulties. Special similarity calculations, as in the case of sequence and structural alignment for macrobiomolecules,^9,10^ are fundamentally challenging tasks. Furthermore, the whole comparison process need to be started from scratch when new batch of records come in. These limitations severely hinders efficient utility and treatment of many types of continuously expanding High Dimensional Data (HDD) with Strong Nonlinear Correlations (SNC). Essentially, the overwhelming majority of interesting data types (e.g. macromolecular structures and sequences, texts, pictures, audio and video) are HDD with SNC. Therefore, it is highly desirable to develop clustering algorithms that do not involve record comparison. We develop such an algorithm in this work and demonstrate its utility in clustering of protein structural domains (PSDs). The major arena of present artificial intelligence research, classification tasks of many types of HDD with SNC (pictures/images, audio and video), engenders labor, time and capital intensive task of human labeling. Essentially, classification is clustering with labels. Therefore, efficient clustering algorithm for HDD with SNC may facilitate classification as well.

### Protein structure comparison and clustering/classification

Protein structure comparison is an important task in computational structural biology. Pairwise comparison for structure similarity is itself a prohibitive task with computational cost being exponential with chain length.^9^ Many excellent approximate/heuristic algorithms^11^ have been developed with great success to overcome this hurdle. However, hierarchical clustering of multiple structure is much more challenging. One way is to perform a progressive approximate multiple structure alignment/comparison.^12^ Apart from inherent approximations, one additional issue of the progressive procedure is that the whole process has to be performed from scratch as new structures are added. A different strategy is adopted by CATH classification,^13^ where partition is first performed according to HMM(hidden markov model) profiles^14^ to establish superfamilies, further agglomerative clustering is performed for upper level topology and architecture and divisive clustering is performed for functional families. This process apparently is subject to limitation of sequence profiling. Both methodologies are not linearly scalable and involves many human decisions.

## Results

### One way encoding based on data type specific unknown SNC

The difficulty of HDD with SNC is mainly the disentangling of inter-dimensional correlations, brute force calculation of which inevitably requires excessively large data sets and prohibitive computational cost. However, clustering of such records does not necessarily demand full understanding of these complex correlations. Since correlations reduce effective degrees of freedom (DOF)/dimension, all HDD with SNC, as a matter of fact, map to manifolds of significantly lower dimension (LD). While it is extremely challenging to directly understand and specify such manifolds, realizing their existence may help us tremendously. As long as composition for a high dimensional (HD) record is given with certain necessary constraints, the inherent SNC will drive it to proper position in the unknown LD manifold. A simple example is that despite numerous possible ways for a given set of students to sit in a given classroom, most probably they sit in a small number of configurations, which would be further restricted to a unique configuration with some additional neighboring constraints without understanding of interactions among students. Therefore, composition of a HD record, plus some additional constraints, may be utilized to encode it for the purpose of placing it correctly in its LD manifold. One step further, composition of a specific data type may be defined to include necessary constraints. Accomplishment of this idea would produce data type specific one-way encoding schemes significantly more efficient than regular encoding (e.g. compression algorithms) that need to be reversed later on and applicable for arbitrary data. Such encoding utilizes unknown data type specific SNC and therefore is irreversible and has to be constructed on a case by case basis. Assume we knew a viable definition of composition for such one way encoding of a given data type, the next key step is to design a bookkeeping strategy so as to cut its LD manifold into segments and consequently realize clustering. In this way, SNC become our tool instead of problem. This idea is illustrated in Fig. 1. It is usually quite difficult to invent a perfect encoding as shown in Fig. 1(c). However, some slight misplacement in LD manifold (Fig. 1(d)) may be good enough for approximate clustering. Of course, an encoding with weak/no constraining power would generate nearly random placement of records in LD manifold as shown in Fig. 1(e). Designing a good encoding is a brain racking task.

**Figure 1:**
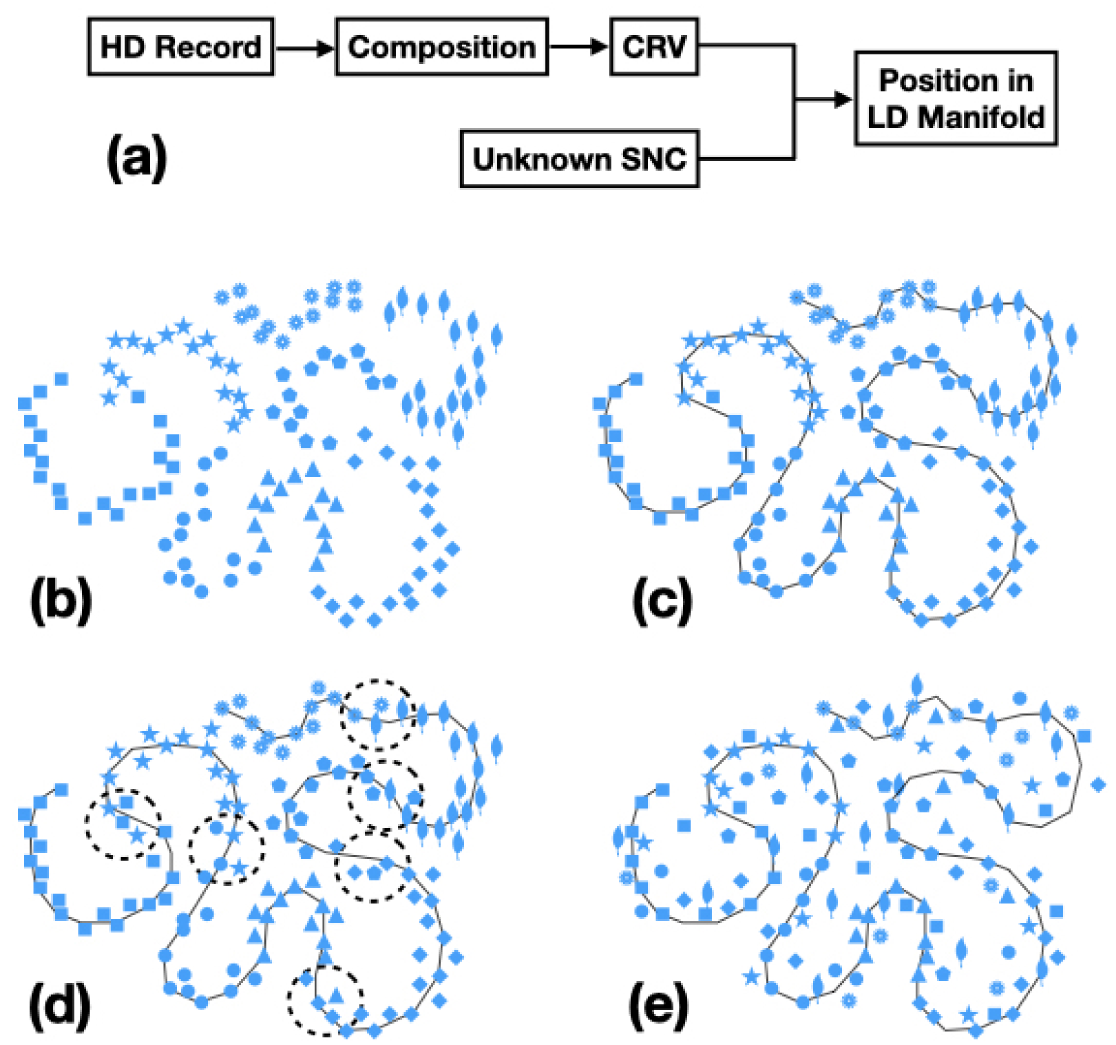
Schematic representation of mapping from nominal high dimensional space to lower dimensional manifold driven by unknown SNC. (a) CRV encoding illustration. (b) scatter record points in high dimensional space. (c) a perfect mapping to manifold (indicated by a thin line). (d) approximate mapping to manifold (misplaced records shown in dashed circles). (e) A failed mapping.

### Composition rank vector tree

Our intuition is that property of a record should be determined by its compositions, and composition of the largest fraction should be of the most importance. Based on this thought we come up with composition rank vector (CRV) tree clustering. There might well be many better encoding based clustering algorithms to be discovered. The CRV tree algorithm is shown below:

1. Define composition for a given data type. Initialize an empty CRV tree (root).
2. For each record:
  a. Count the occurrence for each composition to obtain the CRV with the most populous composition being the first element, the second most populous composition being the second element, etc.
  b. Insert the record into the CRV tree with its path specified by the CRV and the root.
3. Each node and its descendants form a cluster, the CRV tree is a hierarchical clustering. The maximum possible number of nodes for a data type with *n* defined compositions is 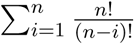, with 0! being 1.
4. For a more quantitative characterization, segments overlap/distance in LD manifold may be evaluated by examining possible local rank reversals of each record in the tree. For example, for a data type with three compositions (*A, B, C, D*). Record 1 (*R*1) has 8 *A*s, 7 *B*s, 4 *C*s and 1 *D* with CRV [*ABCD*], record 2 has 7 *A*s, 8 *B*s and 2*C*s with CRV [*BAC*], these two records have local rank reversals of the first two compositions. This operation is linearly scalable as number of possible reversals to exam do not increase with number of records.

In strong contrast to all present clustering algorithms, no record comparison is involved in this procedure. Consequently, addition of new records doe not interact with existing records in a CRV tree, and the process is linearly scalable to arbitrarily large data sets. We further demonstrate the utility of the algorithm in clustering of PSDs.

### Construction and statistics of PSD CRV tree

We tested three different definition of composition (correspond to three different encodings termed type-I, type-II and type-III respectively) shown below: i) 8 DSSP secondary structure states and gaps are defined as 9 compositions; ii) pair-wise permutation of the 9 type-I compositions are defined as 81 compositions; iii) Combination of 20 regular amino acid and ‘*X*′ with three secondary structure states *H*(helical), *B*(sheet) and *−*(coil), plus a gap denoted as ‘′s, are defined as the 64 compositions (Fig. 2(a)). The first two definitions of composition do not embody the fundamental interaction preferences in proteins, and consequently are not useful in posit PSDs correctly in LD manifold (results not shown). We focus on the clustering results from type-III composition. “depth” and “level” are used interchangeably to describe tree depth.

**Figure 2:**
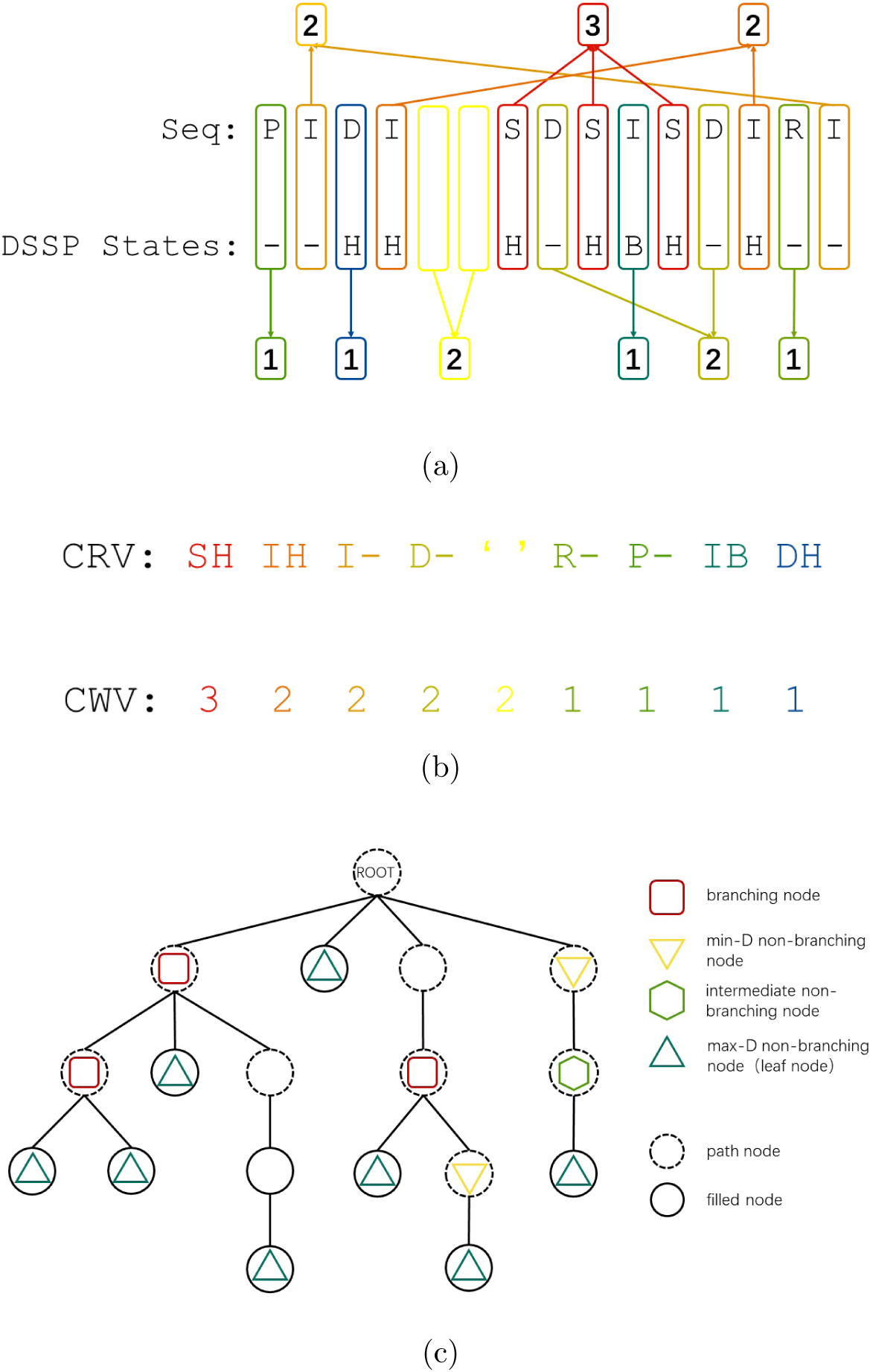
Illustration of CRV construction and CRV tree terminology. (a) Counting the number of different comprising unit according to type-III composition definition for a hypothetical PSD with sequence “PIDI SDSISDIRI”, “H”, “B” and “-” are helix, sheet and coil states respectively. “_” in caption and empty yellow boxes in figure (a) represent discontinuous segments in primary sequence, which are quite often seen in PSDs defined in CATH database. (b) The resulting CRV and CWV (composition weight vector). (c) Terminology for the nodes in a CRV tree. A node does not store any PSD is termed “path node” and “root” is always a path node. Nodes store one or more PSDs are termed “filled nodes”. A node has more than one children node is termed “branching node”, A node has less than two children is termed “non-branching”. Non-branching nodes can be of minimum-depth (min-D), maximum-depth (max-D, or leaf) or intermediate depth (intermediate).

The construction of a CRV (Fig. 2(a),(b)) and terminology of CRV tree nodes (Fig. 2(c)) are illustrated in Fig. 2. We constructed the PSD CRV tree for 427574 PSDs downloaded from CATH database. Details of the PSD CRV tree statistics are listed in the supplementary excel file. Tree branching occurs mainly at depths from 2 to 20 (Fig. 3(a)(b)). The number of minimum-depth (min-D) non-branching nodes decrease sharply from depth level 5 (48262) to 15 (7172) (see supplementary table for detailed information). However, most of PSDs have CRV lengths of approximately 25 - 58, as shown in Fig. 3(c), suggesting nodes in shallow levels of the CRV tree are mainly path nodes. The majority filled nodes have only 1 PSD while significant number of filled nodes have more than one PSDs (Fig. 3(d)). However, this observation does not necessarily indicate the failure of our encoding, it is simply due to the fact that there are many very similar PSDs in the database. We performed sequence alignment for PSDs sharing the same nodes, and find that qualitatively, sequences in deeper level nodes have lower sequence identity (Fig. 3e) but the lowest number is 82%. This may seem counter intuitive as more shared depth in a tree should correspond to more similar structures. However, CRV is only a qualitative description with potentially many different CWVs sharing the same CRV. The longer a CRV is, the more opportunity are available for variation of corresponding CWVs, which is directly associated with sequences given our definition of composition include full resolution for different amino acids.

**Figure 3:**
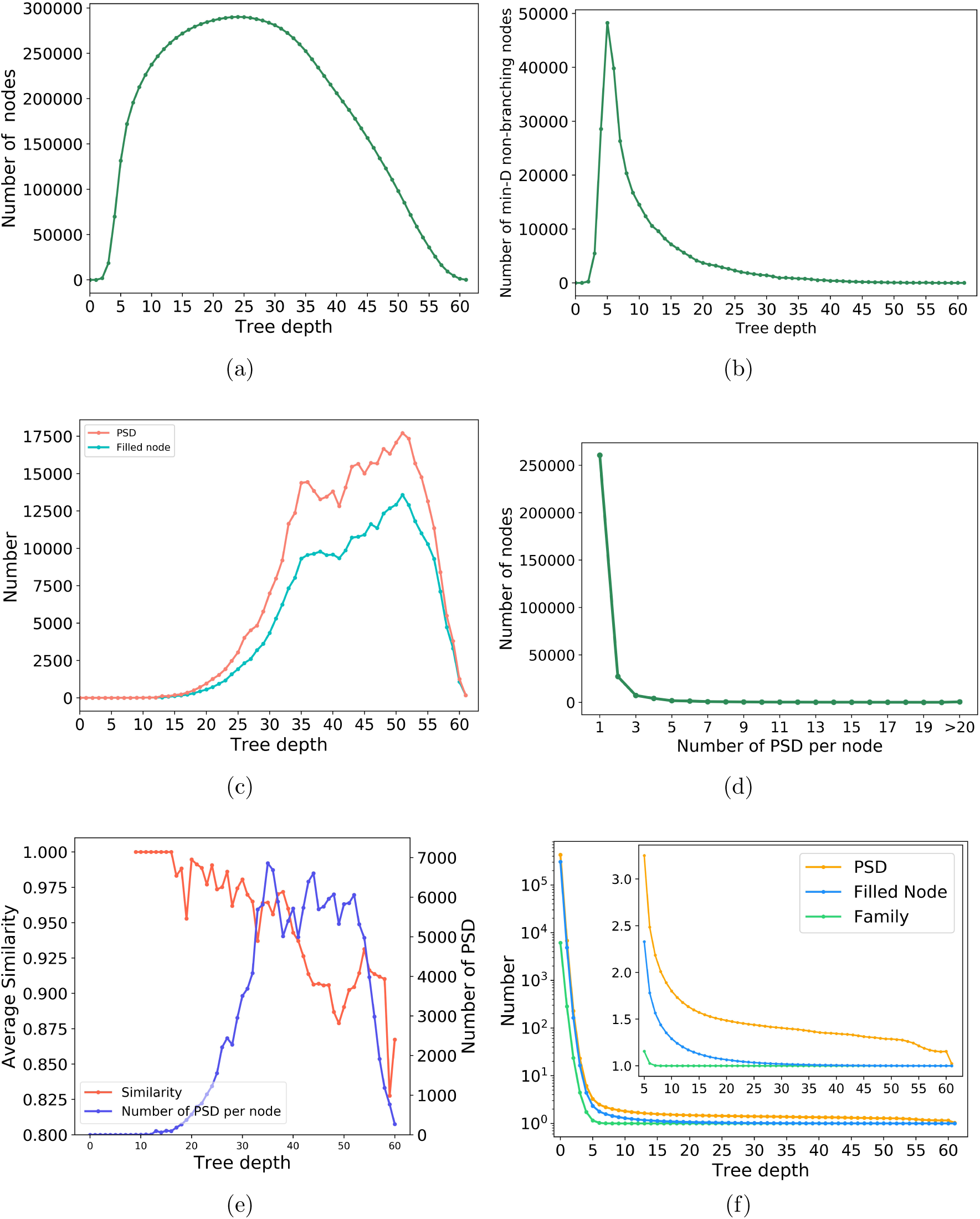
Summary of the PSD CRV tree. (a) Total number of nodes at each level of tree depth. (b) Total number of min-D non-branching nodes at various tree depth. (c) Total number of PSDs and filled nodes at various tree depth. (d) Number of nodes that store various number of PSDs. (e) Average sequence similarity for PSDs sharing same nodes (orange line is for sequence similarity and blue line is for total number of PSDs from those filled nodes with more than one PSD.). (f) Average number of PSDs for subtrees at various depth, inset is in linear scale.

To find a similar PSD for an enquiry PSD in the CRV tree, all one need to do is insert the enquiry PSD in the CRV tree, first look for any PSD in the same node. If none is available, then trace up for the first branching node, PSDs in this ancestor node and all other descendants of which are the most similar ones. This qualitative algorithm does not specify the similarity extent of sibling nodes. One may trace further up in the CRV tree for the second, third branching nodes for further cousins.

### Comparison with the CATH database

Superfamilies are the most important and the first step of PSD clustering by CATH methodology based on sophisticated sequence profiles. Further agglomerative algorithm are utilized to establish upper level clustering. The latest CATH-Plus version 4.2.0 we downloaded has 6119 superfamilies. The 64-composition CRV tree has 63 top-level subtrees (the root is always an empty node) (see supplementary file for details), 1877 second level nodes and 18535 third level nodes. Therefore, some superfamilies are clearly partitioned at this level. Of course, further partition occurs as we go deeper in the CRV tree. We listed all pairs of PSDs and presented the probability of such pairs locate in different CATH superfamilies based on their shared depth and their distances in the CRV tree (Fig. 4). Essentially all PSDs have shared depth larger than 7 belong to the same superfamily. However, there are a few exceptions as indicated in Fig. 4(b). We looked into details of the 5 indicated PSD pairs. Three pairs have both high level of sequence and structural similarity (98% and 99% sequence identity), two pairs have high level of structural similarity by very low level of sequence identity(11% and 13%), see supplementary document for details. These cases suggest that our extremely simple clustering can reveal various neglected similarity by sophisticated CATH procedures.

**Figure 4:**
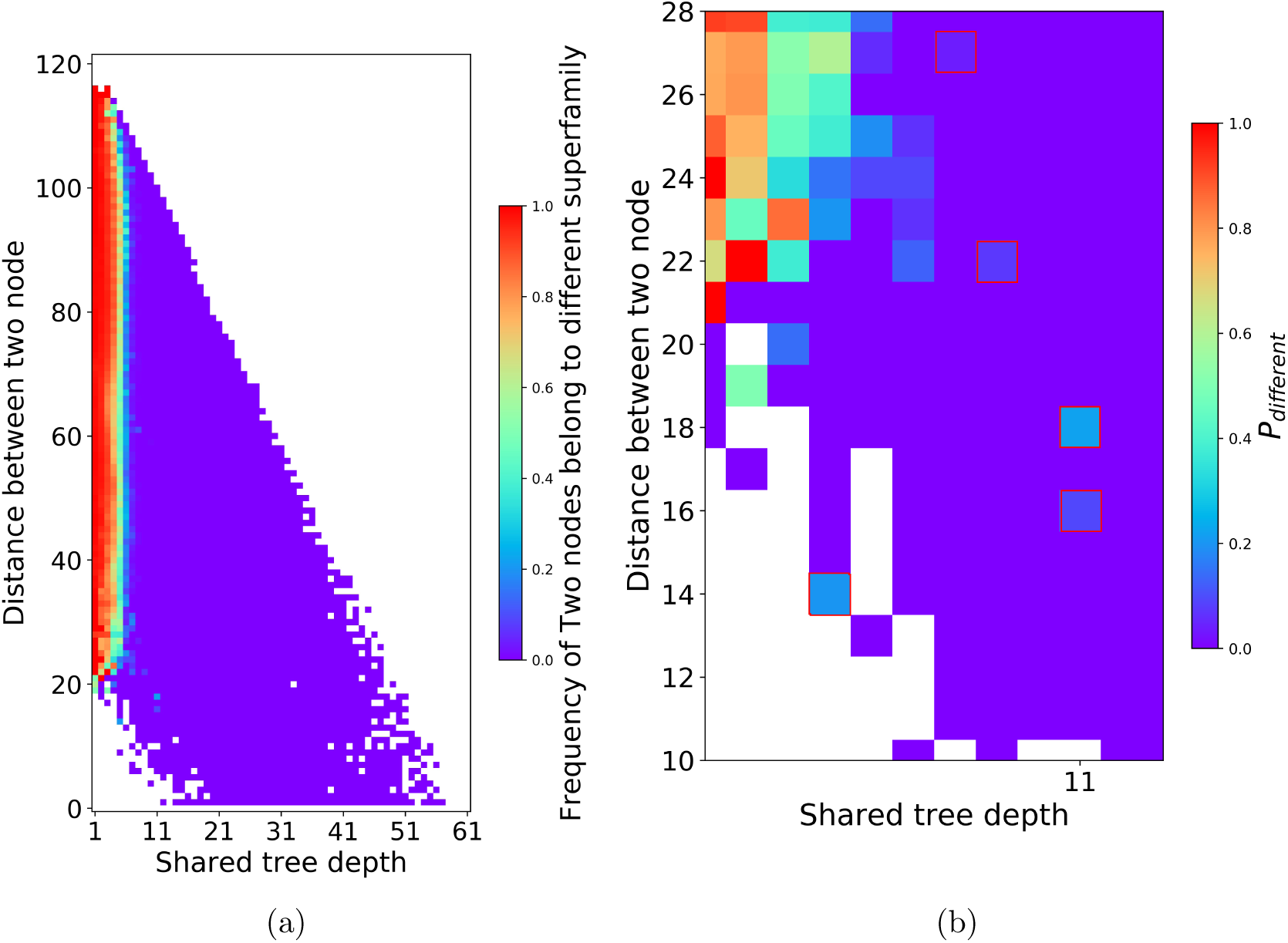
Comparison with CATH superfamilies. For all PSD pairs that have the same shared depth and distance in CRV tree, we calculated the probability that a pair belong to different superfamily as defined by CATH. (a) The full plot. (b) Magnification of the lower left part, with five cases denoted red squares, where PSD pairs with shared tree depth equal to or larger than 7 but belong to different CATH superfamilies, see supplementary materials for details.

## Discussions

As is reflected in the case of PSD clustering, definition of composition is the simple most important factor in CRV tree clustering algorithm. The constraints provided by definition of composition, plus all unknown data type specific SNCs, should be sufficient to posit essentially all records to proper positions in their LD manifold. One necessary consideration for the definition of composition is the different interaction preference among them. Physically, all HDD with SNC have well-defined (yet unknown to us) interaction preferences among its comprising units. Our definition of composition need to capture them to be useful. At one limit, if all defined compositions interact with weak preference, as in our case of 8 DSSP secondary structural states as compositions of PSDs, the corresponding CRVs have weak constraining power, consequently records would not be posited at right location in the corresponding LD manifold and encoding fails (results not shown). This is as expected since most amino acid (AA) can adopt any secondary structural states (though with different extent of preferences), both local and long-range residue preferences are quite weak when only secondary states are in consideration. With both AA identity and secondary structural states, essentially all PSDs are clustered quite successfully as compared with CATH. For any given data type, defining of composition require understanding of their interactions. Therefore, no general recipe is available except necessity of sufficiently strong interaction preference, which need case by case understanding of data type. The fact that no record comparison is needed makes CRV tree clustering very suitable to continuously expanding HDD sets, which essentially include the overwhelming majority of interesting data. More importantly, substitution of record comparison by encoding provides linear time and space computational complexity while realizing hierarchical clustering, this unparalleled scalability makes it an ideal choice for arbitrarily large data sets.

Direct mapping from HDD with SNC to its corresponding LD manifold is an extremely challenging task. Brute force realization of this goal demands exponentially large data sets and exponentially increasing computational costs as a function of dimensionality and this simple fact is widely termed as “the curse of dimensionality”. CRV tree clustering starts with the motivation of utilizing complex unknown mapping between HD record and its position in LD manifold. However, possibility exist that as sufficiently large data set is provided, the target LD manifold will naturally emerge, as relative closeness of different clusters may be characterized by step iv) of the algorithm. While it is difficult to estimate the necessary amount of data a priori, the “sufficiently large” data set should scale exponentially to the dimensionality of corresponding LD manifold instead of original dimensionality of HD record. The dimensionality gap between HD record and corresponding LD manifold is apparently a function of inter-dimension correlations and may well make the difference between the computable and the incomputable.

CRV is apparently not the only one-way encoding of information. Essentially most machine learning algorithms are one-way encoding processes that encode input information to a value as in regression, a class label as in classification, and a number of clusters as in comparison based clustering. For example, neural networks are a very diversified class of trainable one-way encoder. These two types of one-way encoding have fundamentally different motivations. In CRV encoding, one specify some constraints by definition of composition and the derived CRV, then let inherent SNC in data drive the record to proper position in unknown LD manifold. In present machine learning algorithms, the output dimensionality is explicitly specified, and one seeks to find an explicit mathematical map between HD input information and its position in LD label space through repetitive training. Unfortunately, to find a good definition of composition one has to rack his/her brain and test in a laborious way. Therefore making searching of better composition definition trainable is highly desired and this is one planned direction of our algorithmic research. Additionally, development of ensemble methods based on different CRV trees is another potentially productive direction to explore. More importantly, CRV is just one example of utilizing unknown SNC to encode and cluster. Searching better ways of utilizing SNC is a potentially rewarding new direction in machine learning.

Utility of CRV, or any other yet-to-be-discovered encoding based on unknown SNC, is certainly not limited to clustering. It may be utilized to encode complex input information (e.g. picture, texts, video and audio) into vectors and feed into neural networks to combine their respective strength. CRV is a qualitative representation of its corresponding record. In constructing of CRV, composition weight vector (CWV) has to be generated as an intermediate quantitative information. While not utilized in this work, CWV is likely to be useful to further improve CRV tree clustering, as a matter of fact, optional step 4) of the CRV tree clustering actually needs CWV. While CRV can be quite long when a large number of comprising compositions are defined for a given data type, it is likely that only a small number of principle compositions should be sufficient to perform necessary clustering. As demonstrated in case of PSDs, with 64 possible compositions, all CRV tree branching essentially occurs within 10 principal compositions. This could be of great importance for application of CRV tree clustering in more complex data types as pictures, video, audio and texts where number of compositions can be potentially quite large. CRV is a relatively robust way of encoding as small variations (noises) in the most populous compositions usually do not change their ranks.

## Conclusions

In summary, we propose the idea of utilizing unknown SNC to carry our concise data type specific encoding of HDD. CRV tree based clustering algorithm is invented as a specific example of utilizing such encoding schemes. This is the first linearly scalable hierarchical clustering algorithm. Human decisions are limited to definition of composition and all remaining process is rule based with no parameters involved. We demonstrate the successful utility of this algorithm with hierarchical clustering of PSDs. CWVs accompanying CRVs are also important information, while not utilized in this specific case, is necessary for quantitative characterization of cluster distances in manifolds. It is important to note that CRV tree clustering is more a complementary than a competitor of present clustering algorithms due to the fact that it is not useful for both low dimensional data and HDD with weak nonlinear correlations, for which many present algorithms excel. Nonetheless, CRV tree clustering is a general algorithm applicable for HDD with SNC in many subjects beyond biology. This algorithm might be combined with deep learning in a wide variety of classification tasks. We expect more encoding based clustering algorithm to be discovered in the future.

## Acknowledgement

This research was supported by National Natural Science Foundation of China under grant number 31270758

## Notes

### Competing Interest Statement

The authors have declared no competing interest.

